# Structural basis of translation in transcription-translation coupling

**DOI:** 10.1101/2024.03.13.584796

**Authors:** Jing Zhang, Guoliang Lu, Wei Zhou, Vadim Molodtsov, Ziyun Cheng, Yu Zhang, Ruoxuan Li, Zhaoxuan Jiang, Mingxia Yang, Li Li, Huihui Shao, Wei Lin, Xiaogen Zhou, Richard H. Ebright, Jinzhong Lin, Chengyuan Wang

## Abstract

In the bacterium *Escherichia coli*, efficient expression of protein-coding genes is achieved through physical coupling of RNA polymerase (RNAP) synthesizing an mRNA and the first ribosome translating the mRNA, with transcription elongation factors NusG and NusA bridging RNAP and ribosome in a transcription-translation complex (TTC). Recent structural studies suggest that the flexibility in NusG and NusA accommodates the changes in ribosome conformation that occur during the translation cycle, but details have been unclear. Here, we report cryo-EM structures of TTCs at each stage of the translation cycle as an actively translating ribosome approaches to a halted transcription complex. We observe sequential conversion of a loosely coupled TTC (TTC-LC) to a tightly coupled TTC (TTC-B) to a collided TTC (TTC-A) as the ribosome approaches the transcription complex. We show that flexibility of the NusG linker and NusA “pantograph” enable TTC-LC and TTC-B to accommodate ribosome conformation at each stage of the translation cycle. We show that, in TTC-A, steric clash between ribosome and RNAP upon ribosome 30S head swiveling inhibits 30S head swiveling, consistent with translation slowdown, and applies mechanical force to the trailing edge of RNAP, causing transcription termination. Our results define the structures and properties of loosely coupled, tightly coupled, and collided TTCs at each stage of the translation cycle.

## Introduction

In bacteria, all cellular processes take place in the same compartment enabling the first translating (leading) ribosome to coordinate the rate of mRNA translation with the rate of transcription by RNA polymerase (RNAP) for more efficient gene expression^1–13^. This coordination occurs through the formation of transcription-translation coupling complex (TTC)^14–19^. Structural studies of *Escherichia coli* TTCs have defined three classes of potentially functionally relevant TTC: the “loosely coupled TTC” (also referred to as the “long-range coupled TTC”; TTC-LC)^18, 20^, the “tightly coupled TTC” (TTC-B)^15–17, 19^, and the “collided TTC” (TTC-A)^14–17, 19^ (Fig. 1a, Supplemental Fig. 1).

**Fig. 1.**
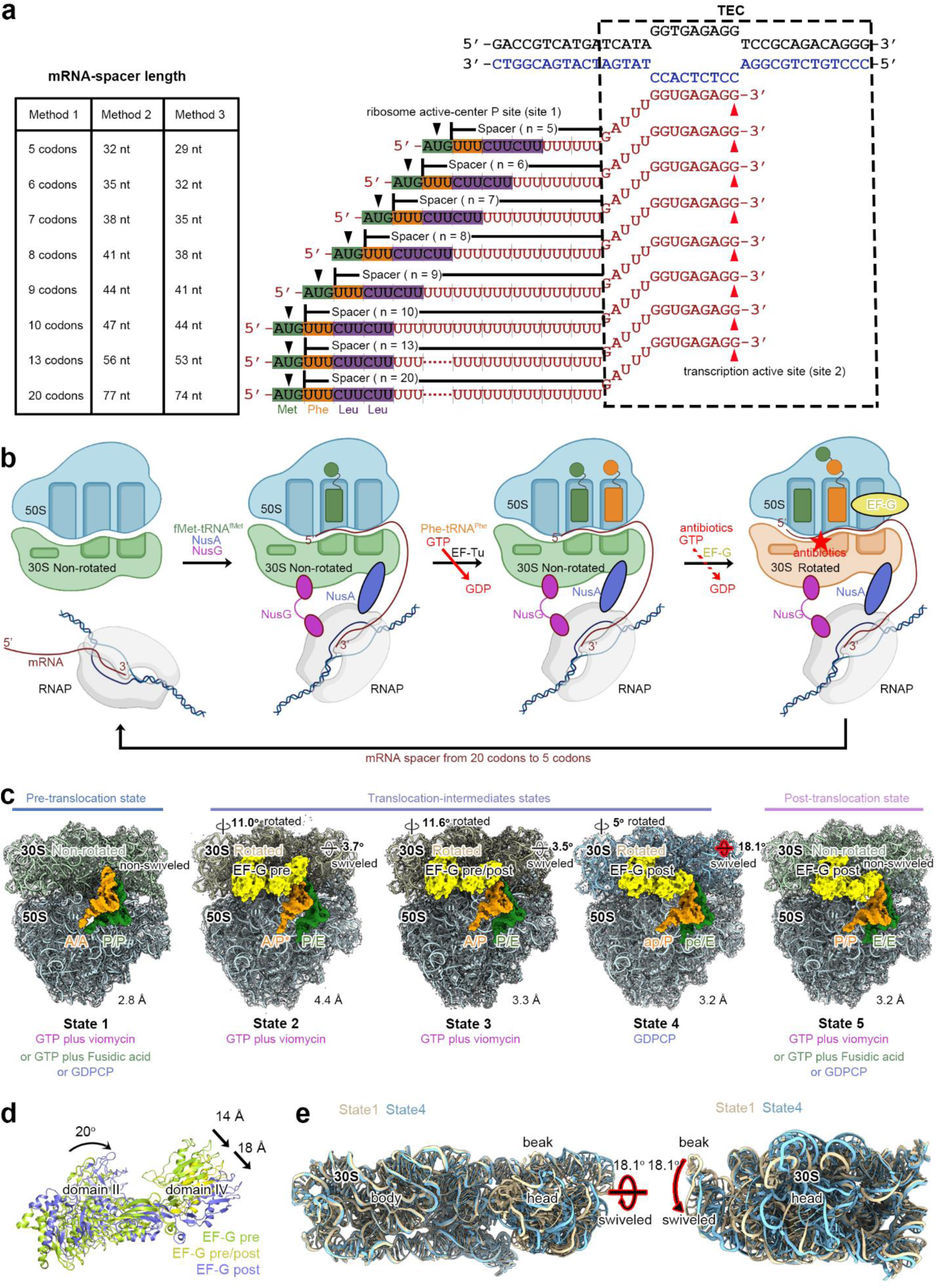
Assembly and structure determination of TTCs at each stage of the translation cycle. **(A)** Left panel, Different representation methods for mRNA-spacer length: Method 1 is from this article, Method 2 is from the Weixlbaumer research group, and Method 3 is from the Zenkin research group; right panel, Nucleic-acid scaffolds, each scaffold comprises non-template- (black) and template-strand (blue) oligodeoxyribonucleotides and one of eight oligoribonucleotides (brick-red), with mRNA-spacer lengthd n of 5, 6, 7, 8, 9, 10, 13, or 20 codons, corresponding to mRNA. Dashed black box labeled “TEC” represents a portion of nucleic-acid scaffold that forms TEC upon the addition of RNAP (9 nt non-template- and template-strand ssDNA segments forming “transcription bubble”, a 9 nt of mRNA engaged with the template-strand DNA as RNA-DNA “hybrid”, and 5 nt of mRNA, on diagonal, in the RNAP RNA-exit channel). The dark green mRNA AUG codon labeled “Met” intended to occupy ribosome active-center P site upon addition of ribosome and tRNA^fMet^: the orange mRNA segment labeled “Phe” represents UUU codon intended to occupy ribosome active-center A site upon addition of tRNA^Phe^: the purple mRNA segment labeled “Leu” represents CUU codons. The “spacer” denotes mRNA-spacer length, in codons, between the initiation codon AUG and the TEC. **(B)** Stepwise reconstitution of translating TTCs. RNAP is first incubated with nucleic-acid scaffold to form TEC (panel 1). TEC is next incubated with 70S ribosome, NusG, NusA, and tRNA^fMet^ to form TTC with tRNA^fMet^ paired with mRNA AUG codon in ribosome active-center P site, and TEC and ribosome bridged by NusG and NusA (panel 2, step 1). TTC is incubated with EF-Tu•tRNA^Phe^•GTP ternary complex to deliver tRNA^Phe^ to ribosome active-center A site (panel 3, step 2). To induce ribosome translocation, TTC is incubated with EF-G•GTP; antibiotics or GDPCP were added at this step if indicated (panel 4, step 3). RNAP is shown in gray, non-template strand DNA, template strand DNA, and RNA are in black, blue, and brick-red, respectively. NusA is shown in light blue, NusG in magenta, tRNA^fMet^ in forest green, tRNA^Phe^ in orange, EF-G in yellow, and antibiotics or GDPCP are depicted as a red star. **(C)** Structures of translation intermediate states of ribosomes showing EM density (grey surface) and fit (ribbons). From left to right, ribosome state 1, resolution 2.8 Å, state 2, resolution 4.4 Å, state 3, resolution 3.3 Å, state 4, resolution 3.2 Å, and state 5, resolution 3.2 Å, respectively. TEC, mRNA, ribosome, NusG, and NusA are shown as ribbon models; tRNAs and EF-G are shown as EM densities. RNAP is shown in gray, non-template strand DNA, template strand DNA, and RNA are in black, blue, and brick-red, respectively. The non-rotated 30S subunit is shown in light green, rotated 30S subunit in wheat, and 50S subunit in cyan. NusA is shown in light blue, NusG in magenta, tRNA^fMet^ in forest green, tRNA^Phe^ in orange and EF-G in yellow, respectively. Rotation angles for the 30S and 50S models are shown at their top-right, with the swivel angle of the 30S head relative to its body annotated on the head. Ribosome stalk is omitted for clarity in this and all subsequent images. **(D)** Structural changes in EF-G during translocation. “EF-G pre” (green), “EF-G pre/post” (yellow) and “EF-G post” (purple) conformations of EF-G are superimposed by structural alignment of 50S subunit. Strait arrows indicate rotational movement of EF-G domain IV between these states. The curved arrow indicates rotation of EF-G body between “EF-G post” and “EF-G pre” or “EF-G pre/post” states. **(E)** Structure comparison of TTC-B state 1 and state 4 by superimposing 30S body. The 16S rRNA in TTC-B is shown as wheat ribbon in state 1 and light blue ribbon in state 4.

In TTC-LC, the ribosome and RNAP are bridged by transcription elongation factor NusG or its paralog RfaH, and optionally further bridged by transcription elongation factor NusA, and the ribosome and RNAP make no or negligible direct interaction with each other (Supplemental Fig. 1a,)^18, 20^. TTC-LC is observed when the mRNA-spacer length between the ribosome active-center P site and RNAP is at least ∼12 codons, accommodates small, ∼1-5 codon, variation in mRNA-spacer length through differences in the extent of compaction of mRNA in the RNAP RNA-exit channel, and accommodates very long, potentially unlimited-length mRNA spacers by looping out of the mRNA between ribosome and RNAP (Supplemental Fig. 1a)^18^, In TTC-LC, the distance between the ribosome mRNA-entry portal and RNAP is ∼80 Å, and the part of the mRNA segment between the ribosome mRNA-entry portal and RNAP closest to the ribosome mRNA-entry portal interacts with the RNA-helicase domain of ribosomal protein uS3 in the ribosome 30S head, and the rest of the mRNA segment between the ribosome mRNA-entry portal and RNAP makes no interactions with ribosome, RNAP, NusG, or NusA, enabling looping out of this part of the mRNA and accommodation of very long mRNA spacers. In TTC-LC, the NusG C-terminal domain interacts with ribosomal protein uS10 in the 30S head, and NusA KH-1 domain interacts with the part of the mRNA segment between the ribosome mRNA-entry portal and RNAP closest to the ribosome mRNA-entry portal, contacting the face of that mRNA segment opposite the face contacted by uS3.

In TTC-B, the ribosome and RNAP are bridged by transcription elongation factor NusG or its paralog RfaH and optionally further bridged by transcription elongation factor NusA, and the ribosome and RNAP make direct interactions with each other (Supplemental Fig. 1b)^15–17^. TTC-B is observed when the mRNA-spacer length is ∼7-12 codons^15–17^, and accommodates small, ∼1-5 codon, variation in mRNA-spacer length through differences in the extent of compaction of mRNA in the RNAP RNA-exit channel (Supplemental Fig. 1b)^15–17^. In TTC-B, the distance between the ribosome mRNA-entry portal and RNAP is ∼45 Å, and the mRNA segment between the ribosome mRNA-entry portal and RNAP interacts with the RNA-helicase domain of ribosomal protein uS3 in the ribosome 30S head. In TTC-B, the RNAP β’ zinc binding domain (ZBD) interacts with ribosomal protein uS3 in the ribosome 30S head, the NusG C-terminal domain interacts with ribosomal protein uS10 in the 30S head, and NusA KH-1 domain interacts with ribosomal proteins uS2 and uS5 in the ribosome 30S body.

In TTC-A, the ribosome and RNAP make direct interaction with each other, and are arranged in a manner that is incompatible with bridging by NusG or its paralog RfaH and incompatible with binding and bridging by NusA (Supplemental Fig. 1c)^14–17, 19^. TTC-A is observed when mRNA-spacer length is ∼4-8 codons and accommodates small, ∼1-4 codon, variation in mRNA-spacer length through differences in the extent of compaction of mRNA in the RNAP RNA-exit channel (Supplemental Fig. 1c)^15–17^. In TTC-A, RNAP is positioned directly over the ribosome mRNA-entry portal, the distance between the ribosome mRNA-entry portal and the RNAP RNA-exit channel is just ∼20 Å, and mRNA proceeds directly into the ribosome mRNA entrance channel from the RNAP RNA-exit channel. In TTC-A, the RNAP β subunit flap-tip helix (FTH) interacts with ribosomal protein uS3 in the ribosome 30S head, the RNAP α^I^ subunit C-terminal domain interacts with ribosomal proteins uS3 and uS10 in the ribosome 30S head, and the RNAP β’ subunit zinc binding domain interacts with ribosomal protein uS4 in the ribosome 30S body.

In each translation cycle, the ribosome 30S subunit “rotates” by ∼12° relative to the ribosome 50S and then “back-rotates “; the ribosome 30S head “swivels” by ∼18° relative to the 30S body and then “back-swivels”; the ribosome binds and then releases translation elongation factors and tRNAs; and the ribosome translocates by 1 codon--i.e., by 3 nt--relative to the mRNA (Fig. 1b-d)^21–24^. In order to maintain transcription-translation coupling across the translation cycle, a functional TTC must be able to accommodate the changes in ribosome tertiary structure, ribosome quaternary interactions, and ribosome translocational state that occur across the translation cycle, and must be able to coordinate the 1-codon, 3-nt step size of the ribosome with the 1-nt step size of RNAP.

Based on structural modelling, TTC-LC and TTC-B have been predicted to able to accommodate the changes in ribosome structure, quaternary interactions, and translocational state that occur across the translation cycle, and thus have been predicted to be translationally active^15–18^, and a detailed hypothesis--the “coupling pantograph” hypothesis--has been set forth for how TTC-LC and TTC-B accommodate these changes. In contrast, based on structural modelling, TTC-A has been predicted to be unable or only partly able to accommodate ribosome 30S head swiveling, and thus has been predicted to be translationally inactive or only partly translationally active^15, 16, 19^. Accordingly, it has been proposed that TTC-LC and TTC-B functionally mediate transcription-translation coupling for actively transcribing ribosomes and actively translating RNAP, and that TTC-A is either non-functional, or functional only in special cases when RNAP is paused or irreversibly arrested, serving to push forward a paused RNAP, resulting in transcription rescue, or to push forward an arrested RNAP, resulting in transcription termination^15–18^. Consistent with these proposals, single-molecule fluorescence-resonance energy-transfer results show that TTCs having NusG-dependence, NusA-dependence, and mRNA-spacer lengths matching those in TTC-LC and TTC-B functionally mediate transcription-translation coupling for actively transcribing ribosomes and actively translating RNAP, and that, when ribosomes approach closer to RNAP, yielding mRNA-spacer lengths consistent with those in TTC-A, ribosomes exhibit translational slowdown and accumulate in translation-cycle fully rotated states^20^. Also consistent with these proposals, biochemical results indicate that TTCs mRNA-spacer lengths matching those in TTC-A can push forward a paused RNAP, resulting in transcription rescue, can push forward an irreversibly arrested RNAP, resulting in transcription termination^8, 12, 13^. However, the hypothesis that TTC-LC and TTC-B are incompatible with ribosome 30S head swiveling has been disputed^10^, and, prior to the work in this report, none of hypotheses and proposals in this paragraph have been tested directly.

It has been shown that TTC-LC can transform into TTC-B upon ribosome catch-up (i.e., when a translating ribosome approaches close enough to RNAP to reduce the mRNA-spacer length to 7-12 codons), and it has been shown that TTC-B can transform into TTC-LC upon RNAP run-ahead (i.e., when a transcribing RNAP moves far enough from RNAP to increase the mRNA-spacer length to ≥12 codons)^18^. Accordingly, it has been proposed that TTC-LC is a functional intermediate in assembling and disassembling TTC-B, mediating pre-TTC-B transcription-translation coupling before a ribosome catches up to RNAP, and mediating post-TTC-B transcription-translation coupling after a ribosome stops moving and RNAP continues moving^18^. TTC-LC and TTC-B have different functional properties^18^. TTC-LC, but not TTC-B, is severely defective in RNA-hairpin-dependent transcription termination. Both TTC-B and TTC-LC are severely defective in Rho-dependent transcription termination.

Here, in order to determine how TTC-LC, TTC-B, and TTC-A interconvert, and in order to determine whether and if so how, TTC-LC, TTC-B, and TTC-A accommodate the changes in ribosome structure that occur in each step of the translation cycle, we determined cryo-EM structures of TTCs in which an actively translating ribosome approached a halted transcription elongation complex (TEC). Starting with an mRNA-spacer length expected to yield TTC-LC, proceeding through mRNA-spacer lengths expected to yield TTC-B, and proceeding through mRNA-spacer lengths expected to yield TTC-A. We obtained 36 structures, comprising structures of TTC-LC in states spanning the full translation cycle and having mRNA-spacer lengths of 20 and 13 codons; structures of TTC-B in states spanning the full translation cycle and having mRNA-spacer lengths of 13, 10, 9, 8, and 7 codons; structures of TTC-A in states of the translation cycle prior to full ribosome 30S swiveling and having an mRNA-spacer length of 7 codons; and structures of RNAP-free translation complexes, resulting from transcription termination, for mRNA-spacer lengths of 6 and 5 codons.

### Assembly and structure determination of TTCs at each stage of the translation cycle

To prepare translating TTCs at each stage of the translation cycle, we used synthetic nucleic-acid scaffolds that contained i) DNA and mRNA determinants that direct formation of transcription elongation complexes (TECs) upon addition of RNAP, ii) an mRNA translation start codon (AUG) that directs formation of translation complex upon addition of 70S ribosomes and tRNA^fMet^, iii) an mRNA phenylalanine codon (UUU), followed by two leucine codons (CUU-CUU), immediately downstream of the translation start codon that enables 1 codon of ribosome translocation upon addition of EF-Tu•Phe-tRNA^Phe^•GTP ternary complex, and iv) mRNA spacers having lengths, *n*, of 20, 13, 10, 9, 8, 7, 6, and 5 codons between (i) and (ii) (Fig 1a). For each nucleic-acid scaffold, we equilibrated nucleic-acid scaffolds with RNAP to form a TEC; then added NusG, NusA, and 70S ribosome and tRNA^fMet^ to form a TTC having the translation start codon and tRNA^fMet^ positioned in the ribosome active-center P site; then added EF-Tu•Phe-tRNA^Phe^•GTP ternary complex to deliver tRNA^Phe^ to the ribosome active-center A site; and, optionally, translation elongation factor EF-G and the non-hydrolyzable GTP analog GDPCP^25^, or EF-G and GTP in the presence of viomycin^26^ or fusidic acid^27^, which arrest the ribosome at specific stages of translocation (Fig. 1b). For each resulting complex, we then determined the structure using single-particle reconstruction cryo-EM (Table S1).

Our procedure enabled trapping of five distinct states corresponding to five distinct stages of the translation cycle (states 1, 2, 3, 4, and 5; Fig. 1c). State 1--obtained using GDPCP, GTP plus viomycin, or GTP plus fusidic acid--corresponds to the non-rotated (0° rotation), non-swiveled (0° swiveling) translation-cycle pre-translocation state, with tRNAs bound in A/A and P/P mode^23, 24^, and with no EF-G present (Fig. 1c). State 2--obtained using GTP plus viomycin--corresponds to the fully rotated (∼11° rotation), partly swiveled (∼4° swiveling) translation-cycle translocation-intermediate state, with tRNAs bound in ribosome A/P* and P/E sites^23, 24^, and with EF-G in a pre-translocated conformation^23, 24^ (Fig. 1c-d). State 3--also obtained using GTP plus viomycin--corresponds to the fully rotated (∼12° rotation), partly swiveled (∼4° swiveling) translation-cycle translocation-intermediate state, with tRNAs bound in ribosome A/P and P/E sites^23, 24^, and with EF-G in a pre-/post-translocated conformation^23, 24^ (Fig. 1c-d). State 4-- obtained using GDPCP --corresponds to the partly rotated (∼5° rotation), fully swiveled (∼18° swiveling) translation-cycle translocation-intermediate state, with tRNAs bound in ribosome ap/P and Pe/E sites^23, 24^, and with EF-G in a post-translocated conformation (Fig. 1c-e). State 5--obtained using GDPCP, GTP plus viomycin, or GTP plus fusidic acid--corresponds to the non-rotated (0° rotation), non-swiveled (0° swiveling) translation-cycle translocation-intermediate state, with tRNAs bound in ribosome an P/P and E/E sites, and with EF-G in a post-translocation conformation^23, 24^(Fig. 1c-d). These Five states, together with stepwise rotation of EF-G domain II--which ultimately positions EF-G domain IV into the ribosome A site (Fig. 1d)--collectively capture the full spectrum of structural dynamics in a translating ribosome.

Finally, the samples obtained were used for single-particle cryo-EM data collection and processing. The collected data were processed using a mask-based 3D cryo-EM classification strategy with multiple rounds employing different mask positions. (Supplemental Figs. 2-8 and Methods). Following the completion of conventional 2D classification, ribosomal particles were subjected to an initial round of 3D classification. In this round, a 3D mask encompassing the entire ribosome was applied, which facilitated the selection of a set of ribosomal complexes from aggregates. The selected ribosomal particles were then processed through a second round of 3D classification, this time using a mask covering the 30S subunit, enabling the separation of ribosomes in distinct translation states. Subsequently, each class representing a different translation state was subjected to further 3D classification using a mask targeting the RNAP region. This step allowed for the isolation of intact transcription–translation coupling complexes from free ribosomes. This classification procedure was iterated until the number of particles corresponding to transcription–translation coupling complexes no longer changed. Ultimately, we obtained 36 structurally distinct TTC classes, encompassing the entire spectrum of translation elongation intermediates and providing detailed snapshots of the translation process within the TTC context (Fig. 2).

**Fig. 2.**
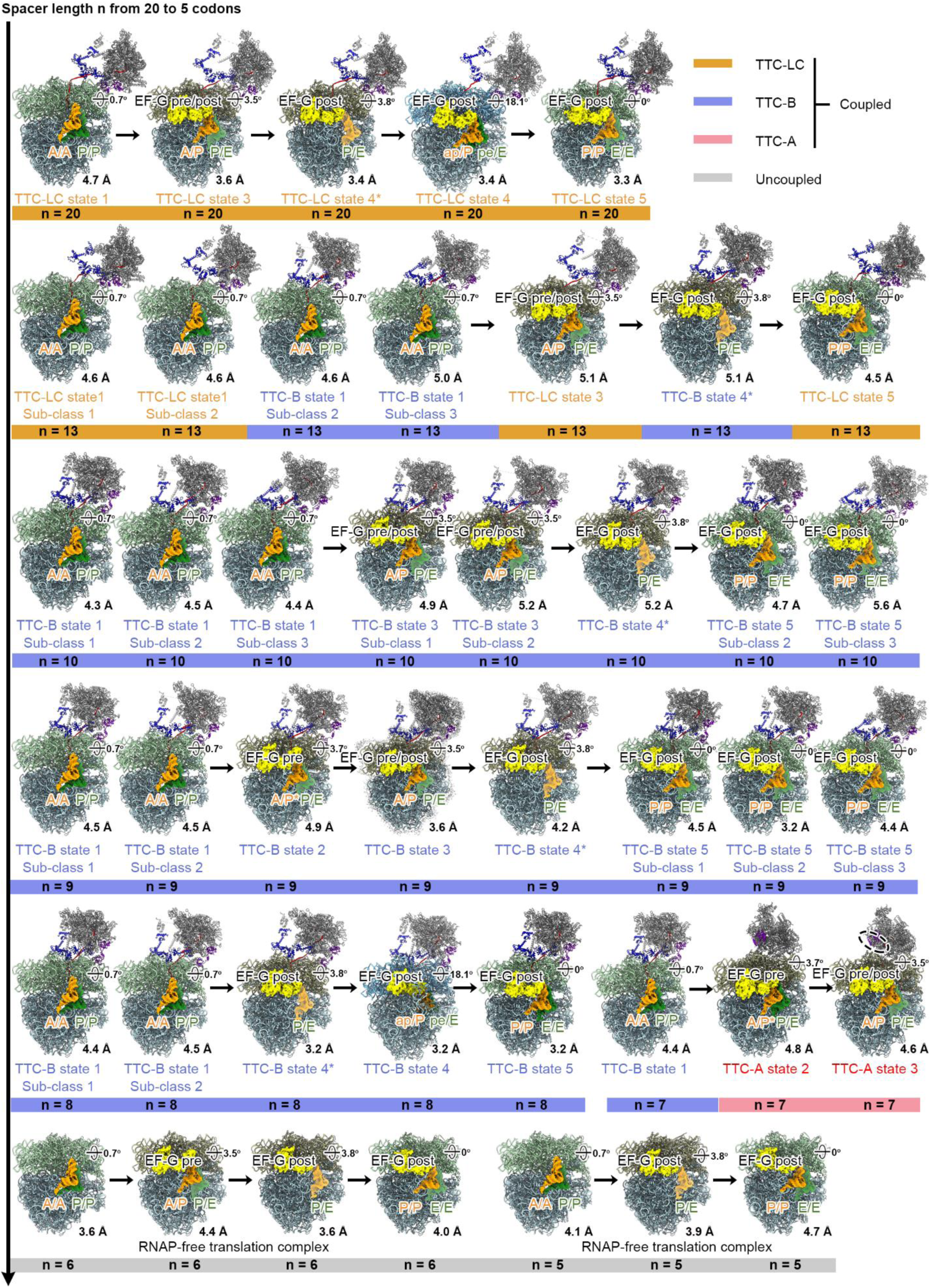
Structures of TTCs at each stage of the translation cycle. Translation elongation cycle in TTCs with different mRNA-spacer lengths. TEC, mRNA, ribosome, NusG, and NusA are shown as ribbon models; tRNAs and EF-G are shown as EM densities. RNAP is shown in gray, non-template strand DNA, template strand DNA, and RNA are in black, blue, and brick-red, respectively. The non-rotated 30S subunit is shown in light green, rotated 30S subunit in wheat, and 50S subunit in cyan. NusA is shown in light blue, NusG in magenta, tRNA^fMet^ in forest green, tRNA^Phe^ in orange and EF-G in yellow, respectively. Rotation angles for the 30S and 50S models are shown at their top-right, with the swivel angle of the 30S head relative to its body annotated on the head. Ribosome stalk is omitted for clarity in this and all subsequent images.

### Structures of TTCs at each stage of the translation cycle

The structures of translating TTC intermediates were determined at resolutions ranging from 3.2 to 5.2 Å, sufficient for model building and consistent with the resolutions of TTCs in recent studies^14–19^ (Fig. 2, Supplemental Figs. 2-8, Table S1). The distinct conformations defined mRNA spacer lengths enabled an accurate reconstruction of the sequential coupling steps. The flowchart in Fig. 2 classifies stable translating TTC intermediates progressing from TTC-LC to TTC-B to TTC-A, and further to transcription termination as mRNA-spacer length shortens from 20 to 13, 10, 9, 8, 7, 6 and 5 codons (Fig. 2).

Long mRNA spacers define the TTC-LC conformation -- it is the sole TTC class when the mRNA-spacer length is 20 codons (Supplemental Fig. 5), and is still present, co-existing with TTC-B, when the mRNA spacer length reduces to 13 codons (Supplemental Fig. 6). TTC-LC is compatible and presents all intermediate states of translating ribosome (Fig. 2). TTC-LC completely disappears from the sample when the mRNA-spacer length reaches 10 codons (Supplemental Fig. 7), and TTC-B becomes the only conformation present with mRNA-spacers ranging from 10 to 8 codons. Similar to TTC-LC, TTC-B is also fully compatible with all intermediate states of the translating ribosome (Fig. 2). When the mRNA-spacer length is further shortened to 7 codons, TTC-B is observed only if the ribosome is in the pre-translocated state. Progression of translation from pre-translocated to translocation-intermediate states results in a collision of the ribosome with RNAP, and conversion of TTC-B to TTC-A (Fig. 2, Supplemental Fig. 8). In this conformation, the ribosome is unable to complete the elongation step, and its translocation ends at states 2 or 3 (Fig. 2). Shorter mRNA-spacer lengths (n = 6 or 5 codons) failed to produce stable TTCs (Supplemental Fig. 9), suggesting that transcription-translation coupling is sterically incompatible under these conditions.

We conclude that mRNA-spacer length determines TTCs class in actively translating TTCs, triggering conversions of TTC-LC into TTC-B at an mRNA-spacer length of ∼13 codons, and conversion of TTC-B into TTC-A at an mRNA-spacer length of ∼7 codons.

### TTC-LC can accommodate each stage of the translation cycle with greater dynamic flexibility

We obtained 9 structures of translating TTC-LC (Figs. 2, 3). The structures have mRNA-spacer lengths of 20 and 13 codons, and define states spanning the full translation cycle, including the translation-cycle pre-translocated state with non-rotated 30S and non-swiveled 30S head (TTC-LC state 1), translation-cycle translocation-intermediate state with rotated 30S and partly swiveled 30S head (TTC-LC state 3), translation-cycle translocation-intermediate state with partly rotated 30S and fully swiveled 30S head (TTC-LC state 4), and translation-cycle post-translocation state with non-rotated 30S and non-swiveled 30S head (TTC-LC state 5) (Figs. 2, 3). For TTC-LC state 1, we obtained two distinct subclasses of translating TTC-LC: subclass 1 and subclass 2 (Fig. 2 and Supplemental Fig. 6). The two subclasses differ from each other by a ∼15° difference in the orientation of RNAP relative to the ribosome 30S head and a ∼5 Å difference in distance between RNAP and the ribosome mRNA-entry portal (Fig. 2 and Supplemental Fig. 10a, b).

**Fig. 3.**
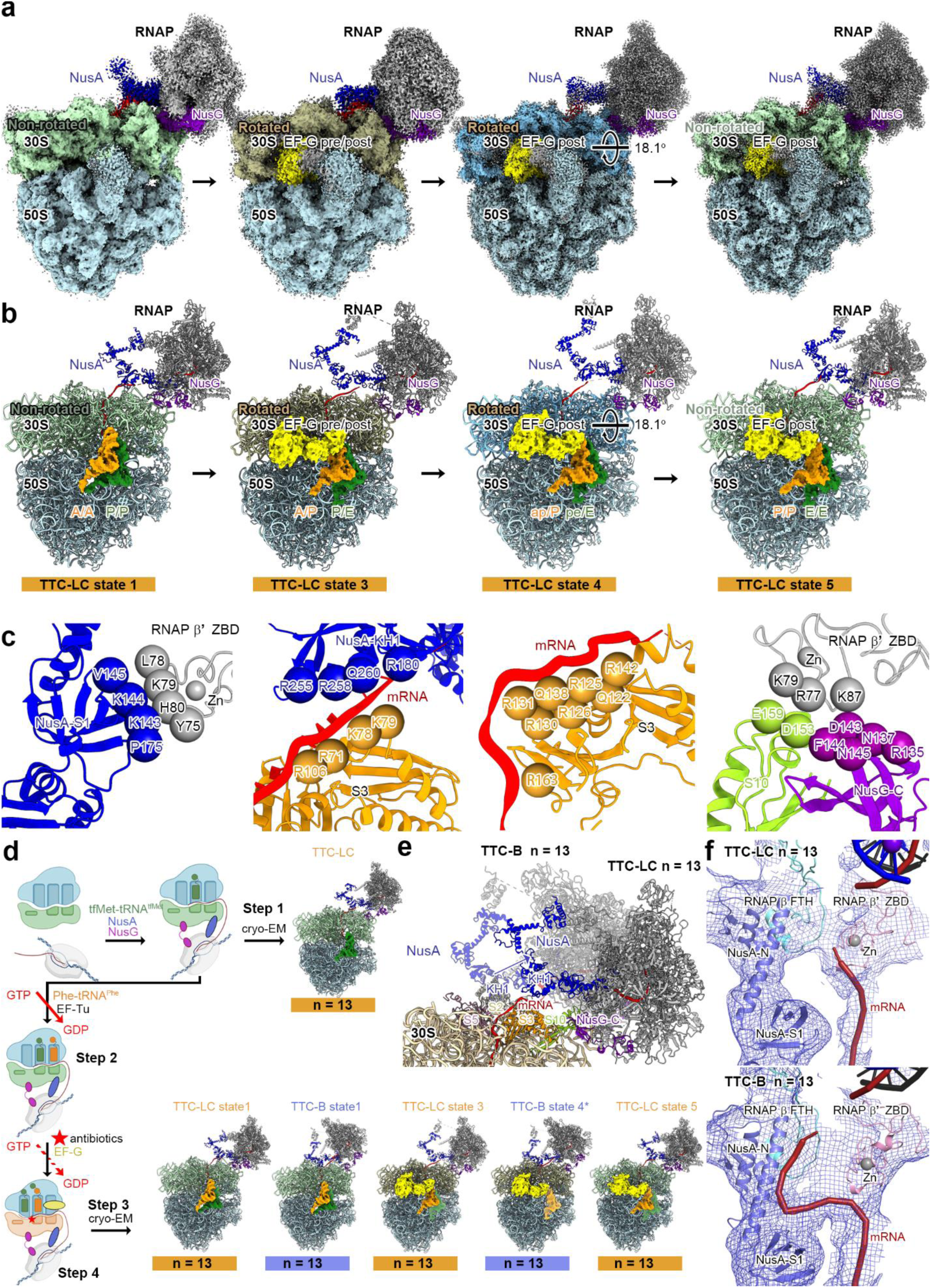
Cryo-EM structures of translating TTC-LC. **(A)** Cryo-EM map of translating TTC-LC. Views and colors as in Figure 2. **(B)** Structure model of translating TTC-LC. Views and colors as in Figure 2. **(C)** Interfaces in TTC-LC. Interactions between RNAP β’ ZBD and NusA-S1 domain (panel 1); interactions between mRNA and NusA-KH1 domain (panel 2); interactions between mRNA and ribosomal protein uS3 (panel 3); interactions between RNAP β’ ZBD, ribosomal protein uS10, and NusG-CTD (panel 4). NusA is shown in blue, β’ ZBD in grey, ribosomal protein uS3 in light orange, mRNA in red, NusG-CTD in purple, ribosomal protein uS10 in light green. **(D)** Schematic diagram of TTC-LC assembly and cryo-EM structures of translating TTC-LC obtained using mRNA with spacer length of 13 codons. Schematic depictions as in Figure 1e; views and colors as in Figure 2. **(E)** Structure comparison of TTC-LC (n = 13 codons) and TTC-B (n = 13 codons) by superimposing 30S body. 16S rRNA in TTC-LC is shown as wheat ribbon. Colors as in Figure 2; TTC-B (n = 13 codons) is shown in semi-transparent colors. **(F)** Comparison of mRNA pathways in TTC-LC (n = 13 codons, upper panel) and TTC-B (n = 13 codons, lower panel) with mRNA spacer length of 13 codons. EM density map near the RNAP exit channel is shown as blue mesh; mRNA, NusA-NTD and S1 domains, RNAP β FTH, RNAP β’ ZBD are shown in brick-red, light blue, cyan, and pink, respectively.

Our ability to obtain structures of translating TTC-LC in a fully rotated state (state 3; Fig. 3) establishes that TTC-LC is compatible with 30S rotation, and our ability to obtain a structure of TTC-LC in a 30S non-rotated state for which a fully rotated state is precursor (state 5; Fig. 3) establishes that TTC-LC is compatible with 30S back-rotation. All interactions between the translation machinery and transcription machinery in translating TTC-LC involve the ribosome 30S subunit (Fig 3). Comparison of structures of translating TTC-LC in non-rotated and rotated states indicates that the structural changes associated with 30S rotation and back-rotation occur far from, and do not affect, the interface between the translation machinery and transcription machinery in TTC-LC (Supplemental Fig. 10c).

Our ability to obtain structures of translating TTC-LC in a fully swiveled state (state 4; Fig. 3a-b) establishes that TTC-LC is compatible with 30S head swiveling, and our ability to obtain a structure of TTC-LC in a 30S non-swiveled state for which fully swiveled states are precursors (state 5; Fig. 3a-b) establishes that TTC-LC is compatible with 30S head back-swiveling. Interactions between the translation machinery and transcription machinery in translating TTC-LC involve only the ribosome 30S head and are made by the RNAP zinc binding domain and NusG C-terminal domain (Fig. 3c). Comparison of structures of translating TTC-LC in non-swiveled and swiveled states indicates that the RNAP zinc binding domain and NusG C-terminal domain maintain without change their interactions with the ribosome 30S head, and move in unison with the 30S head during 30S head swiveling and unswiveling, consistent with one previous hypothesis^18^ (Supplemental Fig. 10d-e)

Our ability to obtain structures of translating TTC-LC in the translation-cycle post-translocated state (state 5; Fig. 3) establishes that translating TTC-LC is compatible with ribosome translocation and the 1-codon decrease in mRNA-spacer length that occurs during ribosome translocation. Comparison of structures of translating TTC-LC in the non-rotated, non swiveled pre-translocated state (state 1) and the non-rotated, non swiveled post-translocated state (state 5) shows no differences in the interface between the translation machinery and the transcription machinery. We infer that translating TTC-LC accommodates ribosome translocation and the 1-codon decrease in mRNA-spacer length that occurs in ribosome translocation through changes in the extent of compaction of mRNA in the RNAP RNA-exit channel, consistent with previous hypotheses^15, 18^ We point out that the existence of TTC-LC subclasses having a ∼15° difference in the orientation of RNAP relative to the ribosome 30S head, and a ∼5 Å difference in the distance between RNAP and the ribosome mRNA-entry portal, potentially helps “buffer” mechanical stresses from ribosome translocation and the concomitant 1-codon reduction in mRNA-spacer length, with transitions from subclass 2 to subclass 1 reducing the distance between RNAP and the ribosome mRNA-entry portal and thereby potentially helping accommodate the 1-codon reduction in mRNA-spacer length.

We conclude that translating TTC-LC can accommodate all changes in ribosome structure, ribosome quaternary interactions, and ribosome translocation states that occur across the translation cycle, and we conclude that TTC-LC is functional in translation, consistent with the proposal^18^ that TTC-LC functions in general transcription-translation coupling in *E. coli*. We further conclude that translating TTC-LC accommodates ribosome 30S head swiveling and back-swiveling by allowing RNAP and NusG C-terminal domain to move in unison with the 30S head.

### TTC-B can accommodate each stage of the translation cycle

We obtained 25 structures of translating TTC-B (Figs. 2, 4). The structures have mRNA-spacer lengths of 10, 9, 8, and 7 codons, and define states spanning the full translation cycle, including the translation-cycle pre-translocated state with non-rotated 30S and non-swiveled 30S head (TTC-B state 1), translation-cycle translocation-intermediate states with fully rotated 30S and partly swiveled 30S head (TTC-B states 2 and 3), translation-cycle translocation-intermediate state with partly rotated 30S and fully swiveled 30S head (TTC-B state 4), and translation-cycle post-translocation state with non-rotated 30S and non-swiveled 30S head (TTC-B state 5) (Figs. 2, 4). For TTC-B state 1, TTC-B state 3, and TTC-B state 5, we obtained distinct subclasses of translating TTC-B: subclass 1, subclass 2, and subclass 3 (Fig. 2 and Supplemental Figs. 2, 3, 4, 7 and 11). The three subclasses differ from each other by stepwise ∼15° differences in the orientation of RNAP relative to the ribosome 30S head, and stepwise ∼5 Å differences in distance between RNAP and the ribosome mRNA-entry portal (Fig. 2 and Supplemental Fig. 11c). In these subclasses, the relative positioning of RNAP and the 30S subunit resembles that observed earlier in the TTC-B subclasses, with the only difference being that the ribosomes in these subclasses are now in distinct translation-cycle states.

**Figure 4.**
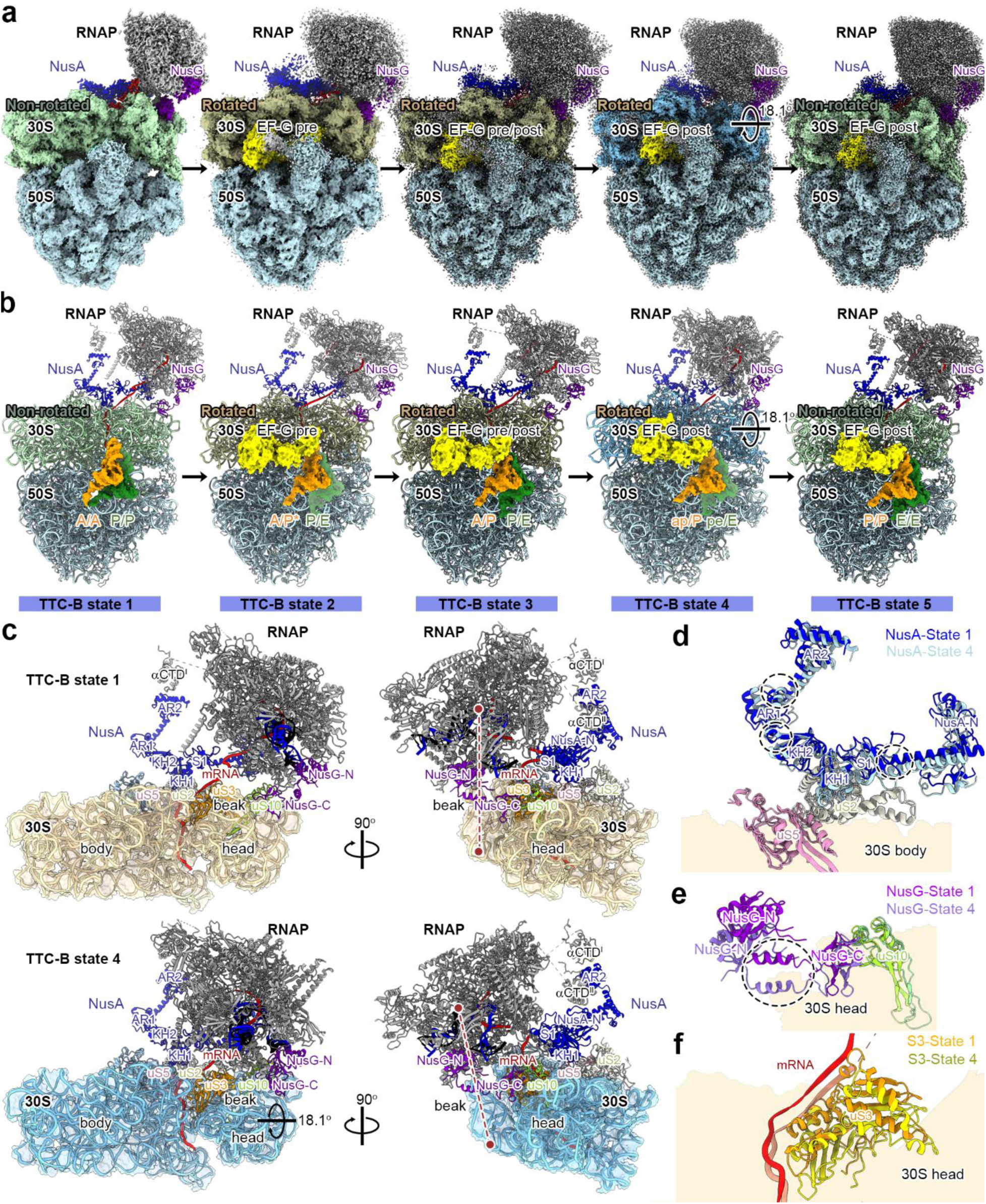
Cryo-EM structures of translating TTC-B. **(A)** Cryo-EM map of translating TTC-B. Views and colors as in Figure 2. **(B)** Structure model of translating TTC-B. Views and colors as in Figure 2. **(C)** Two orthogonal view orientations showing 30S head swiveling in TTC-B state 1 (upper panel) and TTC-B state 4 (lower panel). The 16S rRNA in TTC-B is shown as wheat ribbon in state 1 and light blue ribbon in state 4, mRNA in brick red, ribosomal protein uS2 in brown, ribosomal protein uS3 in orange, ribosomal protein uS5 in pink, ribosomal protein uS10 in light green. In TTC-B state 4 30S head undergoes an 18.1° clockwise rotation as compared to TTC-B state 1 (black arrow). **(D)**, **(E), (F)** Close-up views of NusA, NusG, and mRNA stretching. Flexible linkers in NusA (NTD-S1 linker, AR1-KH2 linker, AR1-AR2 linker; dashed line circle) and flexible linker in NusG (NusG linker; dashed line circle), bridging ribosome and RNAP. Colors as in Figure 2; state 4 is shown in semi-transparent colors.

TTC-LC converts to TTC-B, and indeed, TTC-B makes 37.6% of all TTCs assembled on a nucleic-acid scaffold with the mRNA-spacer length of 13 codons (Fig. 3d). During the structural rearrangement of TTC-LC to TTC-B, RNAP rotates ∼60° relative to the NusG C-terminal domain-uS10 contact, and the KH-1 domain of NusA slides along the mRNA into a shallow pocket on the 30S subunit surface formed by the ribosomal proteins uS2 and uS5, forming a stable interaction with them (Fig. 3e). An additional distributed contact, here termed “cradle”, was observed to form in TTC-B, involving ∼ 9 nt long mRNA segment and positively charged residues of NusA-NTD, RNAP β flap-tip helix, and RNAP β’ zinc binding domain. The “cradle” previously has been observed in structures where NusA interacts with His pause hairpin loops and terminator hairpins^28^ (Fig. 3f and Supplemental Fig. 10f).

Our ability to obtain structures of translating TTC-B in fully rotated states (states 2 and 3; Fig. 2) establishes that TTC-B, like TTC-LC, is compatible with 30S rotation, and our ability to obtain a structure of TTC-B in a 30S non-rotated state for which fully rotated states are precursors (state 5; Fig. 2) establishes that TTC-B, like TTC-LC is compatible with 30S back-rotation. All interactions between the translation machinery and transcription machinery in translating TTC-B involve the ribosome 30S subunit (Fig. 2). Comparison of structures of translating TTC-B in non-rotated and rotated states indicates that the structural changes associated with 30S rotation and back-rotation occur far from, and do not affect, the interface between the translation machinery and transcription machinery in TTC-B (Supplemental Fig. 11a).

Our ability to obtain structures of translating TTC-B in a fully swiveled state (state 4; Fig. 2) establishes that TTC-B, like TTC-LC, is compatible with 30S head swiveling, and our ability to obtain a structure of TTC-B in a non-swiveled state for which fully swiveled states are precursors (state 5; Fig. 2) establishes that TTC-B, like TTC-LC, is compatible with 30S head back-swiveling. Interactions between the translation machinery and transcription machinery in translating TTC-B involve both the 30S head and the 30S body, with the RNAP zinc binding domain and NusG C-terminal domain interacting with the 30S head, and with NusA KH-1 domain interacting with the 30S body (Fig. 4c-f). Comparison of structures of translating TTC-B in non-swiveled and swiveled states indicates that the RNAP zinc binding domain and NusG C-terminal domain maintain without change their interactions with the ribosome 30S head, and move in unison with the 30S head during 30S head swiveling and unswiveling consistent with one previous hypothesis^15, 18^ (Fig 4c, e), and show that the NusA maintains--without change its interactions with the ribosome 30S body and RNAP during 30S head swiveling and unswiveling, accommodating changes in relative orientation of RNAP and 30S body through changes in the conformations of flexible connectors between the NusA N-terminal and S1 domains, KH-1 and AR1 domains, and AR1 and AR2 domains within the NusA “pantograph”^15, 18^, consistent with previous hypotheses (Fig 4d)^15, 18^.Our ability to obtain structures of translating TTC-B in the translation-cycle post-translocated state (state 5; Fig. 2) establishes that translating TTC-B, like TTC-LC, is compatible with ribosome translocation and the 1-codon decrease in mRNA-spacer length that occurs during ribosome translocation. Comparison of structures of translating TTC-B in the non-rotated, non-swiveled pre-translocated state (state 1) and the non-rotated, non-swiveled post-translocated state (state 5) shows no differences in the interface between the translation machinery and the transcription machinery. We infer that translating TTC-B, like translating TTC-LC, accommodates ribosome translocation and the 1-codon decrease in mRNA-spacer length that occurs in ribosome translocation through changes in the extent of compaction of mRNA in the RNAP RNA-exit channel, consistent with previous hypotheses^15, 18^. We point out that the existence of TTC-B subclasses having stepwise ∼15° increases in the rotational orientation of RNAP relative to the ribosome 30S head, and stepwise ∼5 Å increases in distance between RNAP and the ribosome mRNA-entry portal, like existence of TTC-LC subclasses, potentially helps “buffer” mechanical stresses from ribosome translocation and the concomitant 1-codon reduction in mRNA-spacer length, with transitions from subclass 3 to subclass 2 and from subclass 2 to subclass 1 reducing the distance between RNAP and the ribosome mRNA-entry portal and therby potentially helping accommodate the 1-codon reduction in mRNA-spacer length.

We conclude that translating TTC-B can accommodate all changes in ribosome structure, ribosome quaternary interactions, and ribosome translocation states that occur across the translation cycle, and we conclude that TTC-B is functional in translation, consistent with the proposal^15, 18^ that TTC-B mediates general transcription-translation coupling in *E. coli*. We further conclude that translating TTC-B accommodates ribosome 30S head swiveling and back-swiveling by allowing RNAP and NusG C-terminal domain to move in unison with the 30S head, and by exploiting flexing of connector elements within the NusA “pantograph”^15^ to compensate for changes in the orientation and distance of RNAP and the ribosome 30S body.

### TTC-A is incompatible with major stages of the translation cycle

We obtained 2 structures of translating TTC-A (Figs. 2, 5). The structures have an mRNA-spacer length of 7 codons and define TTC states only for the part of the translation cycle that precedes full ribosome 30S head swiveling: i.e., translation-cycle translocation-intermediate states with rotated 30S and partly swiveled 30S head (TTC-A states 2 and 3) (Figs. 2, 5).

**Figure 5.**
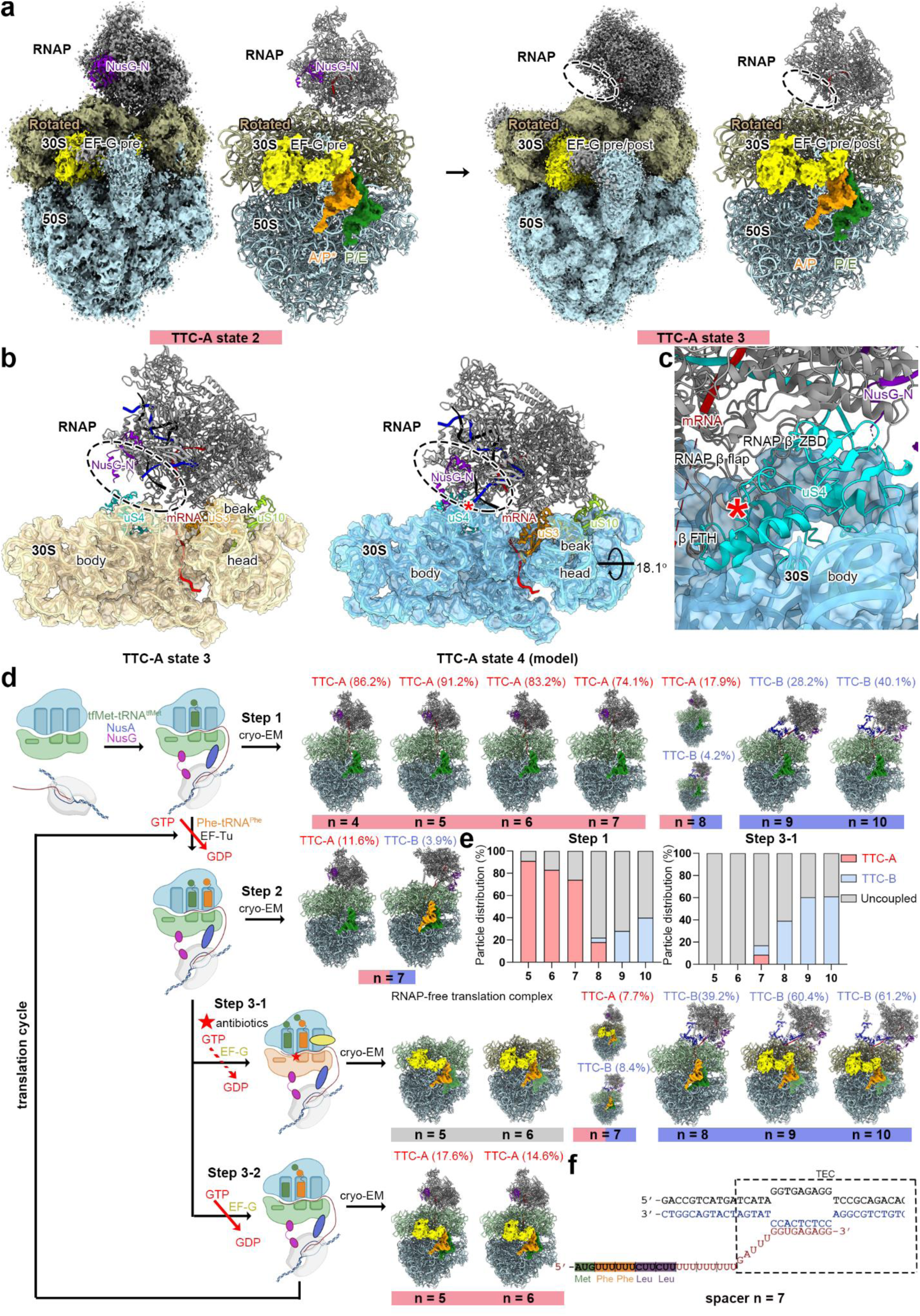
Cryo-EM structures of translating TTC-A. **(A)** Cryo-EM map and structure model of translating TTC-A. Views and colors as in Figure 2. **(B)** Two orthogonal view orientations showing 30S head swiveling in TTC-A state 3 (left panel) and modeled TTC-A state 4 (model, right panel). Views and colors as in Figure 3c. Missing EM density of β’ ZBD, RNAP aCTD^I^ and NusG-NGN domain are indicated by a dashed oval. **(C)** A zoom-in view of structural model of TTC-A in state 4. The red asterisk indicates the location of the clash between ribosomal protein uS4 and the RNAP β flap and β′ ZBD domains. **(D)** Schematic of the sample preparation and cryo-EM structures determination. The workflow is similar to that in Figure 1e, with a modification at step 3, which is split into two parallel reactions: step 3-1 (identical to Figure 1e) involves the addition of EF-G•GTP, antibiotics, or GDPCP to the complex; step 3-2 omits the addition of any antibiotics or GDP analogs. Representative structures are shown in each step, with the corresponding particle distribution data displayed. **(E)** Particle distribution of TTC-A, TTC-B, and RNAP-free translation complex (post-transcription-termination complex) for different mRNA-spacer lengths. Right panel: particle distribution of TTC-A, TTC-B and RNAP-free translation complex (post-transcription-termination complex)in step-1 TTCs with mRNA-spacer lengths of 5, 6, 7, 8, 9, or 10 codons; right panel: distribution of TTC-A, TTC-B and RNAP-free translation complex (post-transcription termiunatoin complex)in step 3-1 TTCs with spacer lengths of 5, 6, 7, 8, 9, or 10 codons. **(F)** Nucleic-acid scaffold with mRNA designed for two rounds of translocation in the presence of tRNA^Phe^.

TTC-B loses its buffering capacity when mRNA-spacer length shortens to a critical threshold of 7 codons, at which point mRNA spans the shortest sterically possible distance between ribosome and RNAP for transcription-translation coupling, and only subclass 1 (the shortest distance from the TEC active site to the ribosome mRNA-entry portal) is observed in the sample. It is likely that in this conformation the ribosome may occasionally directly transmit mechanical force to RNAP as it slows down and fails to maintain a minimum distance.

Our ability to obtain a structure of translating TTC-A in a fully rotated state (state 3; Fig. 2a) establishes that TTC-A is compatible with 30S rotation. All interactions between the translation machinery and transcription machinery in TTC-A involve the ribosome 30S subunit (Fig. 2). Comparison of structures of TTC-A in non-rotated and rotated states indicates that the structural changes associated with 30S rotation occur far from, and do not affect, the interface between the translation machinery and transcription machinery in TTC-A (Supplemental Fig. 12a).

Our inability to obtain a structure of TTC-A in the fully swiveled state (state 4; Fig. 2), together with our inability to obtain a structure of translating TTC-A in the state that follows the fully swiveled state (state 5; Fig. 2) indicate, consistent with prediction^15^, that TTC-A likely is impaired in full 30S head swiveling and impaired in translation. Two additional points support this inference. First, structural modeling of TTC-A in a fully swiveled state (i.e, as state 4) indicates that full swiveling of the 30S head results in severe steric clash between ribosomal protein uS4 in the 30S head and the RNAP β’ zinc binding domain and RNAP β flap-tip helix (Fig. 5b-c). Second, all efforts to obtain translating TTC-A having mRNA-spacer lengths shorter than 7 codons were unsuccessful, yielding, instead, RNAP-free translation complexes produced by transcription termination (Fig. 2; see next section).

We conclude that translating TTC-A can partly accommodate 30S rotation in ribosome structure that occur across the translation cycle, but cannot accommodates ribosome 30S head swiveling and back-swiveling with would yield steric clash between ribosome and RNAP upon ribosome 30S head swiveling, inhibits 30S head swiveling, consistent with translation slowdown, and applies mechanical force to the trailing edge of RNAP, causing transcription termination.

### Late translocation intermediate states trapped by antibiotics trigger TTC-A uncoupling

In our experimental system, TTCs could not be assembled on nucleic-acid scaffolds having mRNA-spacer lengths <7 codons, although TTC-As having shorter mRNA-spacer lengths have been reported previously^14–17, 19^ To address this apparent discrepancy, we conducted a systematic investigation of TTC assembly at different stages, using a nucleic-acid scaffold having an mRNA-spacer length of 7 codons.

We assembled the complexes in the same manner as described above but examined their state by cryo-EM at step 2 (Fig. 5). Our ability to obtain two classes of TTCs—the original TTC-A, with only tRNA^fMet^ positioned in the ribosomal P/P site, indicating successful assembly initiation TTCs (Fig. 5d, Supplemental Fig. 13); and TTC-B, with tRNAs bound in the ribosomal A/A and P/P sites, indicating that EF-Tu has successfully loaded tRNA^Phe^ into the A site—demonstrates the progression of the initiation complex. Additionally, our ability to obtain TTC-B at this step, together with previous reports showing that only TTC-A is present at step 1 when the mRNA spacer length is 7 codons, indicates that a subset of TTC-A complexes is converted into TTC-B during the process of tRNA delivery by EF-Tu. Given that previous studies have demonstrated that the 30S subunit undergoes swiveling and back-swiveling during tRNA delivery by EF-Tu, we propose that in the TTC-A state, the tight interaction between RNAP and the ribosome during 30S swiveling promotes the conversion of a subset of TTC-A complexes into the TTC-B state. Concurrently, only 15.5% of translation complexes contained RNAP at this stage. This significant decrease from the 74.1% observed in step 1 (n = 7 codons) (Fig. 5e) indicates that approximately 80% of the original TTCs have undergone transcription termination. We propose that the swiveling motion also drives transcription termination in TTC-A.

Next, we assembled the complexes in the same manner as described above, except that no antibiotics or GDPCP were added to inhibit ribosome activity, allowing translation to proceed through a complete cycle (Fig. 5d, Supplemental Fig. 14). Our ability to obtain TTC-A with the ribosome captured in the post-translocated state indicates that, in the absence of ribosome inhibition, the complex does not dissociate at the mRNA-spacer length of 6 codons. Repeating the experiment in the same manner and allowing the TTCs to complete two translation cycles (Fig. 5f, Supplemental Fig. 15), we were again able to obtain TTC-A in the post-translocated state. This indicates that, in the absence of ribosome inhibition, the complex does not dissociate at an mRNA-spacer length of 5 codons. By comparing with the results obtained under ribosome-inhibited conditions, we propose that translation-specific inhibitors trap ribosomes in unstable intermediate states, causing them to dissociate under the conformational stress of 30S head swiveling.

Notably, even without antibiotics, the yield of transcription-translation complexes (TTCs) dropped substantially from 74.1% in step 1 to 17.6% in step 3-2 (n from 7 to 6 codons), indicating that the TTC-A state is intrinsically unstable during active translation and susceptible to spontaneous uncoupling. This instability is further evidenced by the mRNA-spacer-length-dependent abundances of TTC-A and TTC-B (Fig. 5e): while TTC-A predominated (up to 90%, n = 5 codons) at step 1 across 5–8 codons, it was only detectable at 7 codons (merely 8% of particles) by step 3-1. This sharp decline, also observed during the TTC-A to TTC-B transition, reflects the low resilience of TTC-A engaged in major conformational changes.At an mRNA-spacer lengths of 7–8 codons, TTC-A and TTC-B maintain a dynamic equilibrium, readily interconverting between states (Fig. 5e).

We conclude that, with mRNA-spacer length of 7 codons, TTC-A state serves as a central intermediate susceptible to two distinct fates during active translation, it can either undergo EF-Tu-mediated conversion to TTC-B coupled with 30S swiveling or promote transcription termination. TTC-A is inherently unstable during active translation, and translation inhibitors trap ribosomes in these unstable swivel-driven intermediates that trigger TTC-A uncoupling and transcription termination.

## Discussion

Our results define the structural basis of translation in transcription-translation coupling. We determined 36 atomic structures that define the interconversion of translating TTC-LC, TTC-B, and TTC-A; define how the changes in ribosome conformation, ribosome quaternary interaction, and ribosome translocational state that occur in the translation cycle are accommodated in translating TTC-LC, TTC-B, and TTC-A; demonstrate that TTC-LC and TTC-B, but not TTC-A, are compatible with all steps of the translation cycle; indicate that formation of TTC-A inhibits ribosome 30S head swiveling; and indicate that formation of TTC-A having an mRNA-spacer length ≤7 codons promotes transcription termination.

Our results show that the flexibility of the bridging interactions made by NusG and NusA in TTC-LC amd TTC-B, particularly the flexibility of the NusA “pantograph,” accommodate changes in ribosome conformation, ribosome quaternary interactions, and ribosome translocational state across the translation cycle. The observed flexibility of the bridging interactions made by NusG and NusA serves the goals of absorbing and dissipating mechanical stresses caused by movements of translating ribosome. Key to this stress-buffering capacity of the bridging interactions by NusG and NusA are the multi-domain structures and domain-specific interactions of NusG and NusA, in which rigid domains bound to ribosome and RNAP are connected by flexible linkers that are unable to efficiently transmit mechanical stresses.

Our results showing that TTC-LC and TTC-B are compatible with all steps in the translation cycle provides strong support for the hypothesis^15, 18^ that TTC-LC and TTC-B are translationally active and that TTC-LC and TTC-B functionally mediate general transcription-translation coupling in *E coli*.

Our results showing that TTC-LC transforms into TTC-B when a translating ribosome advances sufficiently close to RNAP confirms previous evidence for transformation of TTC-LC into TTC-B upon ribosome catch-up^18^ and the proposal that TTC-LC is a functional intermediate in assembling TTC-B, mediating pre-TTC-B transcription-translation coupling before a ribosome catches up to RNAP on ribosome catches up to RNAP^18^.

Our results showing that formation of TTC-A inhibits ribosome 30S head swiveling is consistent with, and account for, results of single-molecule fluorescence resonance energy transfer experiments showing translational slowdown, and accumulation of translation-cycle fully rotated states, as a translating ribosome closely approaches and collides with RNAP^20^.

Our results showing that formation of TTC-A with an m RNA-spacer length <7 codons promotes transcription termination is consistent with, and accounts for, results of biochemical experiments showing transcription termination when a ribosome collides with RNAP^13^. A growing body of evidence indicates that factor-dependent transcription termination, across all domains of life from bacteria to humans, and for most transcription termination factors, involves the application of mechanical force to the TEC trailing edge, specifically to the mouth of the RNAP RNA-exit channel or the RNAP zinc binding domain^29–32^, that results in one or more of (1) forward translocation of the TEC without nucleotide addition (hypertranslocation); (2) extraction of RNA from the TEC (RNA extraction); and (3) reorganization of TEC structure (allostery). Our proposal that ribosome 30S swiveling in TTC-A with an mRNA-spacer length <7 codons causes transcription termination by sterically clashing with, and applying mechanical force to, the transcriptional TEC trailing edge, specifically to the mouth of the RNAP RNA-exit channel and the RNAP zinc binding domain fits this pattern in detail.

After our initial preprint on this work was posted, Mahamid and co-workers posted a preprint on cryo-electron tomagraphy analysis of translating TTCs in *Mycoplasma pneumoniae*^33^. The structural basis of transcription-translation coupling in *M. pneumoniae* is different from in *E. coli* ^19,33^. The orientation of RNAP relative to the ribosome is different in *M. pneumoniae* coupled TTCs, and NusG does not bridge ribosome and RNAP in *M. pneumoniae* coupled TTCs^33^. Nevertheless, Mahamid and co-workers find that *M. pneumoniae* possesses loosely coupled TTCs that are compatible with all stages of the translation cycle, tightly coupled TTCs that are compatible with all stages of the translation cycle, and possesses collided complexes that are inactive, or only partly active, in translation^33^, strikingly analogous to our findings for *E. coli*.

A priority for future studies is the question of whether, and if so how, TTCs coordinate the 1-codon, 3-nt step size of the ribosome with the 1-nt step size of RNAP, structural studies, particularly time-resolved structural studies, with TTCs having both an actively translating ribosome and an actively transcribing RNAP, potentially could address this question.

